# Mimicry embedding for advanced neural network training of 3D biomedical micrographs

**DOI:** 10.1101/820076

**Authors:** Artur Yakimovich, Moona Huttunen, Jerzy Samolej, Barbara Clough, Nagisa Yoshida, Serge Mostowy, Eva Frickel, Jason Mercer

## Abstract

The use of deep neural networks (DNNs) for analysis of complex biomedical images shows great promise but is hampered by a lack of large verified datasets for rapid network evolution. Here we present a novel “mimicry embedding” strategy for rapid application of neural network architecture-based analysis of biomedical imaging datasets. Embedding of a novel biological dataset, such that it mimics a verified dataset, enables efficient deep learning and seamless architecture switching. We apply this strategy across various microbiological phenotypes; from super-resolved viruses to *in vivo* parasitic infections. We demonstrate that mimicry embedding enables efficient and accurate analysis of three-dimensional microscopy datasets. The results suggest that transfer learning from pre-trained network data may be a powerful general strategy for analysis of heterogeneous biomedical imaging datasets.

## Introduction

Artificial Neural Networks (ANN) excel at a plethora of pattern recognition tasks ranging from natural language processing (1) and facial recognition (2) to self-driving vehicles (3, 4). In biology, recent advances in machine learning and Deep Learning (5–7) are revolutionizing genome sequencing alignment (8), chemical synthesis (9, 10) and biomedical image analysis (11–13). In the field of computer vision, convolutional neural networks (CNNs) perform object detection and image classification at a level matching or surpassing human analysts (14). Despite this, CNN-based architectures often poorly recognise unseen or transformed (e.g. rotated) data due to the use of max or average pooling (15). While pooling allows CNNs to generalize heterogenous data, positional information is ignored. This leads to prioritization of smaller image features and results in an inability of the network to “see the big picture”. To circumvent this, dynamically routed capsule-based architectures have been proposed (15, 16). These architectures are nested allowing for the retention of image feature positional information, and optimization of CNN performance on images with a larger field of view.

However, these architectures remain data-hungry and often perform poorly on small biomedical datasets of high complexity (17). One major reason for this is the lack of large, balanced well-verified biological datasets (18), akin to MNIST (19) and ImageNet (20) that allow for rapid algorithm evolution. To circumvent this, ANN analysis of biomedical images can be aided through transfer learning (21, 22). For this, weights of a network trained on one dataset are transferred onto a fully or partially identical untrained network which is then trained on a biomedical dataset of a similar nature (22). This approach shortens training time and is generally considered to be more efficient than random weights initialization strategies (21, 22).

Here, we describe a novel data embedding strategy we term, ‘mimicry embedding’ that allows researchers to circumvent the need for verified biomedical databases to perform ANN analysis. Mimicry embedding involves transforming biomedical datasets such that they mimic verified non-biomedical datasets thereby allowing for mimicry weights transfer from the latter. By embedding 3D, fluorescent image-based vaccinia virus and *Toxoplasma gondii* host-pathogen interaction datasets to mimic grey-scale handwritten digits, we demonstrate that mimicry weights transfer from MNIST (19) allows one to harness the performance of cutting-edge ANN architectures (CapsNet) for the analysis of biomedical data. Furthermore, the high accuracy of the embedded datasets may allow for their use as novel verified biomedical databases.

## Results

More often than not host-pathogen biomedical datasets are not large enough for deep learning. However, we reasoned that advances in high-content fluorescence imaging (23) which allow for 3-D, multi-position single-pathogen resolution can serve to increase the size of datasets for ANN analysis (13). To classify single-pathogen data in 3D biomedical images we developed ‘ZedMate’, an ImageJ-Fiji (24) plugin that uses the Laplacian of Gaussian spot detection engine of TrackMate (25). We challenged ZedMate with multi-channel, 3D fluorescent images of late timepoint vaccinia virus (VACV) infected cells (Fig. 1a and Fig. 1). Owing to its large size, well-defined structure and multiple layers of resident proteins that distinguish different virus forms, VACV has the features needed for complex fluorescence microscopy-based biomedical particle analysis (Fig. 1b). By detecting and linking individual virions within an image across the Z-dimension, ZedMate transforms a series of 2D images into a 3D dataset (Fig. 1b).

**Fig. 1.**
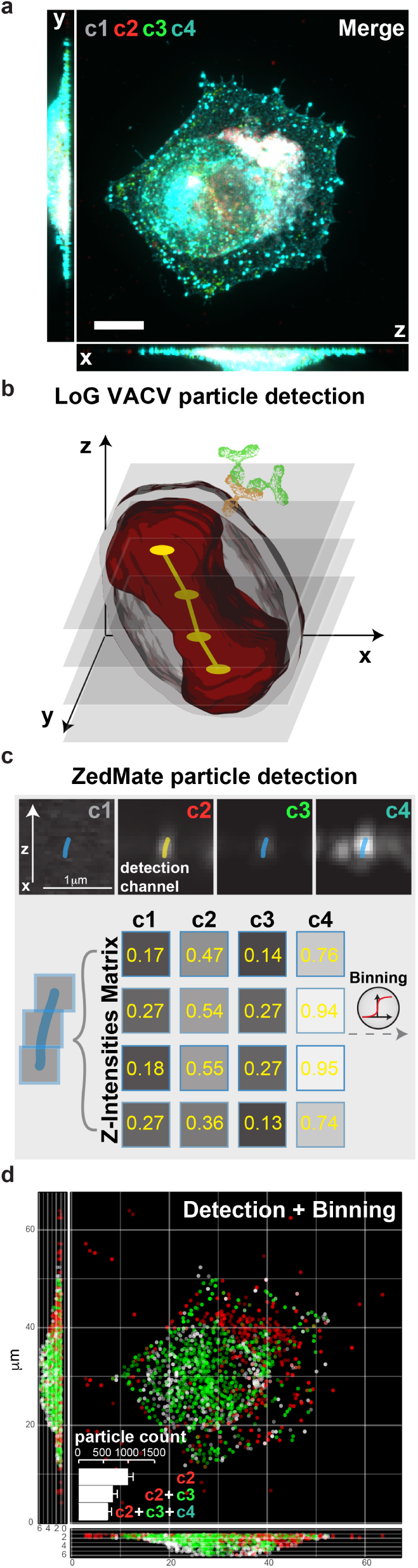
ZedMate facilitates detection and classification of VACV particles in infected cells. (see also Fig. 1). **a**, Merged four channel fluorescent image of a HeLa cell infected with VACV (see Fig. 1a for channel details). Scale bar; 10 µm. **b**, Illustration of Laplacian of Gaussian (LoG)-based VACV particle detection in 3D. Dumbbell shape (red) represents a particle sliced in optical Z-sections (semitransparent grey) providing point signal for LoG detection (yellow) and connected in Z (not to scale). **c**, Intensity measurement from detected particles represented as a Z-profile intensity matrix. **d**, 3D plot of detected particles color-coded according to detected channels and virion category (see Fig. 1b for details). Quantification of different particle types is inset. N=30 cells, error bars; + SEM.

From the original four-fluorescent channel composite ZedMate generates grayscale images that preserve the intensity distribution across the z-dimension of each detected channel (Fig. 1c, upper). From this, fluorescence intensity matrices of each channel per Z-plane are then generated for individual particles (Fig. 1c; lower). Using these matrices and accounting for the 3D positional information of the detected particles, ZedMate reconstructions can be plotted (Fig. 1d). Intensity analysis across all channels allows for binning of virions into three categories consistent with their biological readouts (Fig. 1b).

Initial reconstructions indicated that ZedMate cannot distinguish between incoming cell-free virions and newly replicated cell-associated virions based solely on c1 and c2 intensities (Fig. 1d and S1). To improve the precision of ZedMate-based binning we devised a binary ML/DL strategy relying on manual annotation to separate cell-free from cell-associated virions. To maintain the spatial information acquired in ZedMate we attempted to train the capsule ANN (CapsNet) (15) on this annotated dataset. These initial attempts failed likely due to the small size and complexity of the dataset, two things CapsNet struggles with (17).

To circumvent these issues we decided to harness the state-of-the-art performance of CapsNet on the relatively simple grayscale dataset, MNIST (15). To generate weights matching our binary classification problem, the handwritten digits in MNIST were separated into two classes: <5 and ≤5. With no changes to CapsNet other than restricting its output to two classes, this network converged with 99.6% accuracy (Fig. 2a).

**Fig. 2.**
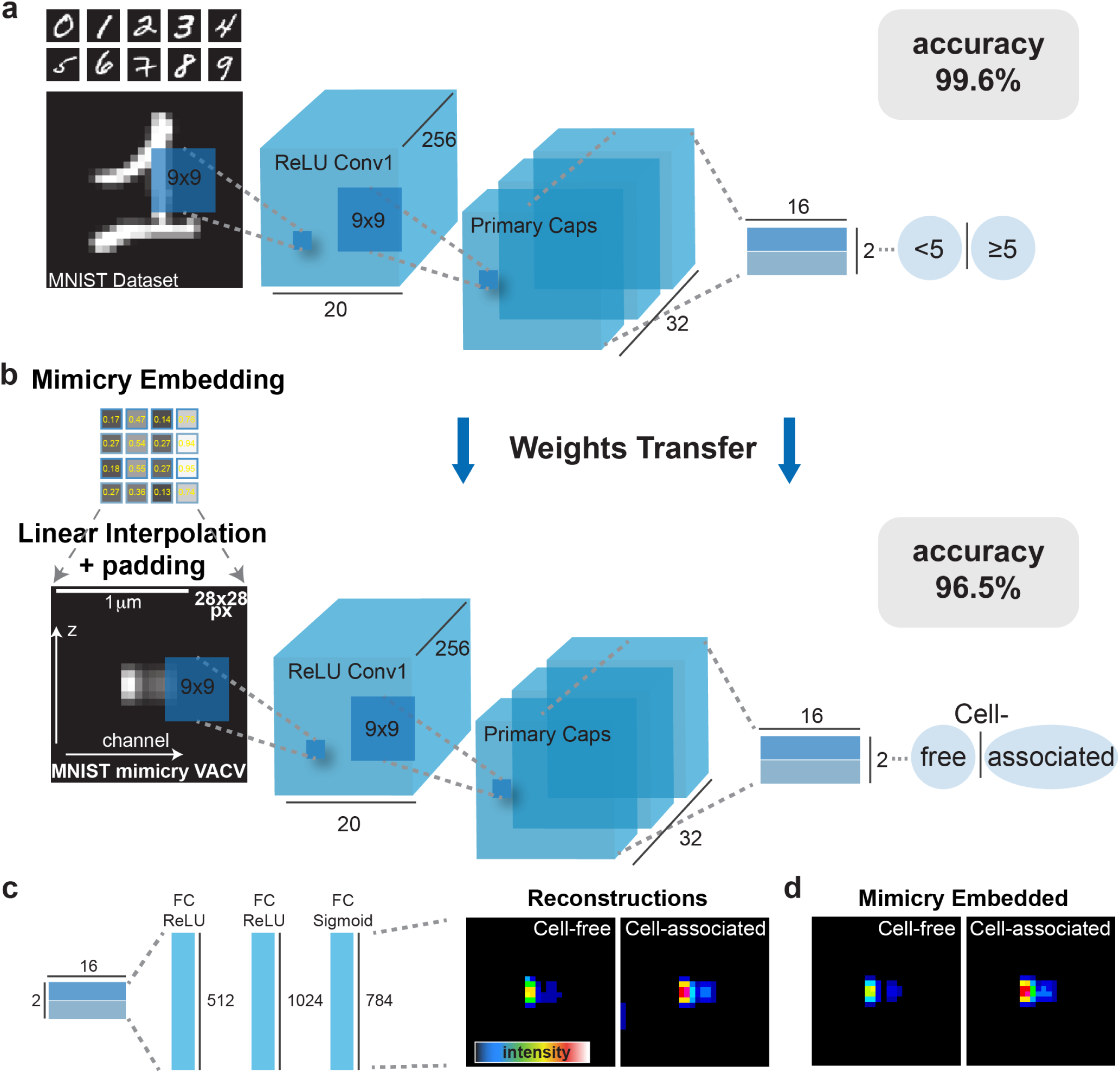
Mimicry embedding allows for separation of cell-free and cell-associated VACV particles through weights transfer from a binary MNIST dataset. **a**, CapsNet architecture for training on the MNIST hand-written digits dataset repurposed into a binary classification problem (<5 or *≤*5) prior to CapsNet weights transfer. **b**, Mimicry embedding of VACV Z-profiles detected by ZedMate. The intensity matrix of fluorescence signal (see Fig. 1) was embedded to mimic MNIST data using linear interpolation and padding Scale bar; 1 µm. CapsNet architecture - with pre-trained weights from a – for training on mimicry embedded VACV particles. **c**, Reconstructed particle profiles of the virions separated as cell-free and cell-associated by CapsNet. **d**, Representative mimicry embedded VACV particles for comparison to c.

To allow for transfer learning from this network to our biomedical dataset we designed a vector embedding strategy we term “mimicry embedding”. For this, the tensors of each virion’s multi-channel, fluorescence Z-profiles from ZedMate are assembled across the X-axis. This is followed by linear interpolation and padding which serve to centre the virion in a 28×28 pixel image such that the resulting data mimics the grayscale MNIST dataset (Fig. 2b). With this approach we aimed to preserve the weights of early CapsNet layers by maintaining the binary MNIST CapsNet architecture and performing weights transfer. Training on our mimicry-embedded real-world dataset achieved 96.5% accuracy (96.2% precision, 96.2% recall) at separating cell-free from cell-associated virions (Fig. 2b and 1a-d for classifier training).

The CapsNet generator was used to visualize how the trained ANN distinguished between cell-free and cell-associated virions with such accuracy. The reconstructions indicated that cell-free virions were elongated with moderate intensity profiles while cell-associated virions were compact and very bright (Fig. 2c). The reconstructions were in agreement with mimicry embedded virions suggesting that these properties yielded the base for the high classification accuracy (Fig. 2d).

To verify this strategy, we performed inference on an unseen (separate from training and validation sets) experimental dataset. Fig. 3 shows the workflow from an input four-channel image (Fig. 3a and S1a,b), to detection and binning of virions (Fig. 3b), followed by mimicry embedding and CapsNet separation of cell-free versus cell-associated virions (Fig. 3c). The results indicate that our model allows for accurate classification of virions into four biologically relevant classes within unseen datasets (Fig. 3d).

**Fig. 3.**
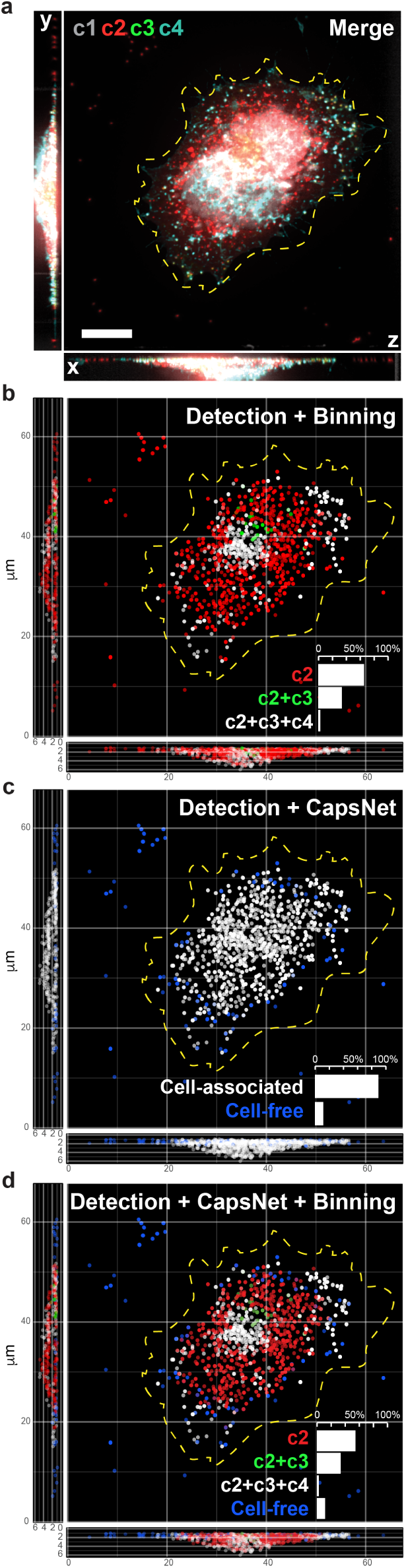
Inference demonstrates that mimicry embedding and trained CapsNet allows for efficient classification of VACV particles into four biological classes. **a**, Merged four channel fluorescent image of a HeLa cell infected with VACV previously unseen by CapsNet (see Fig. 1a for channel details). Scale bar; 10 µm. **b**, Respective ZedMate particle detection and classification by conventional binning of fluorescence intensities. **c**, Respective inference of cell-free and cell-associated particles detected by ZedMate, mimicry embedded and predicted by a trained CapsNet (see Fig. 2b,c). **d**, Combined ZedMate particle detection with mimicry embedded and trained CapsNet results in classification of four types of biologically relevant VACV particles. Inset contains quantification of the particle types in the respective image.

We’ve established that mimicry embedding and weights transfer allows us to distinguish between incoming cell-free and newly assembled cell-associated virions at late time-points after infection. Next, we asked if this approach could also be used to classify extracellular versus intracellular virions during virus entry, a single-cell assay that often requires specific antibodies or labelling strategies and labour-intensive manual annotation. Considering these common limitations, we generated a training dataset that would allow for generalization of this approach. Early infected cells, virions seen in c1, were stained with common fluorescent DNA (c2) and actin (c3) dyes. To circumvent hand-labelling of the training data, immunolabelling to distinguish between intra- and extracellular virus (c4) was used as a weak labelling (26) strategy (Fig. 4a and S3a). After ZedMate detection and transformation of individual particles, intra- and extracellular virus weak labelling (c4) was removed for mimicry embedding. By maintaining our binary MNIST CapsNet architecture and performing weights transfer, we could achieve 82% accuracy (81.3% precision, 81.4% recall) in differentiating between intra- and extracellular virions in the absence of specific-antibody labelling and manual annotation (Fig. 1b-e for classifier training).

**Fig. 4.**
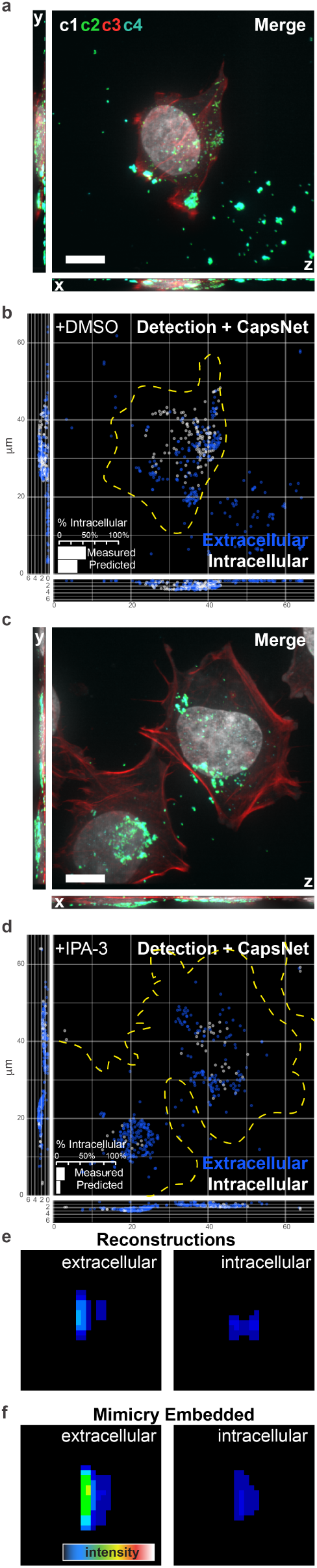
Mimicry embedding can be used for weak-labelling particle classification. **a**, Merged four channel fluorescent image of a HeLa cell infected with VACV previously unseen by CapsNet (see Fig. 1a for channel details). **b**, ZedMate detection and trained CapsNet predicted extracellular and intracellular particles. Quantification of intracellular particles is inset. **c**, Merged four channel image of HeLa cell infected with VACV and treated with the entry inhibitor, IPA 3, previously unseen by CapsNet. **d**, ZedMate detection and trained CapsNet of intracellular and extracellular particles. Quantification of intracellular particles is inset. **e**, Representative reconstruction profiles of extra- and intra-cellular virions. **f**, Representative mimicry embedded extra- and intra-cellular VACV particles for comparison to e. N=40 untreated and treated cells each. Scale bars a-d;10 µm.

To estimate accuracy, inference was performed on an unseen dataset in which intra- and extracellular virions were quantified using c1-c4 (measured)-inclusive of extracellular virion weak labelling – or only c1-c3 (predicted) (Fig. 4b). A 86% match between measured and predicted quantification of intracellular particles was seen (Fig. 4b; inset). This indicates that weak labelling can effectively substitute for manual annotation of training datasets when classifying intra- and extracellular virion signals. As an additional test of the ANN, we generated a dataset skewed for extracellular virions by blocking virus entry with IPA-3 (27, 28) (Fig. 4c). Consistent with its performance (Fig. 1b-e), a 93% match between measured and predicted quantifications of intracellular particles was seen (Fig. 4d). Finally, when we visualized the reconstructions of intra- and extra-cellular virion classes, extracellular virions appeared brighter and more elongated in the Z-direction than intracellular ones (Fig. 4e). This was in agreement with their mimicry embedded counterparts (Fig. 4f), explaining the ANNs ability to accurately predict between and quantify these two virion classes.

To assess the general applicability of our mimicry embedding approach, we acquired a biomedical imaging dataset of cells infected with an EGFP-expressing version of the parasite *Toxoplasma gondii* (*Tg*-EGFP).

While *Tg*-EGFP is readily visualized by conventional microscopy, detecting and quantifying intracellular viability at the single parasite level is challenging (13). To generate a *Tg* viability training dataset, cells infected with *Tg*-EGFP (c1) were fixed and stained with fluorescent markers of DNA (c2), and host cell ubiquitin (c3) which was used a weak label to annotate the subset of “unviable” parasites (13, 29) (Fig. 5a). Individual particle detection and transformation in ZedMate was followed by mimicry embedding in the absence of c3 weak labelling. After weights transfer from the binary MNIST CapsNet architecture (illustrated in Fig. 2b), and fine tuning on the *Tg*-EGFP we achieved 70% accuracy (precision 72.5%, recall 70.7%) in the absence of specific viability labelling (Fig. 1a-d for classifier training).

**Fig. 5.**
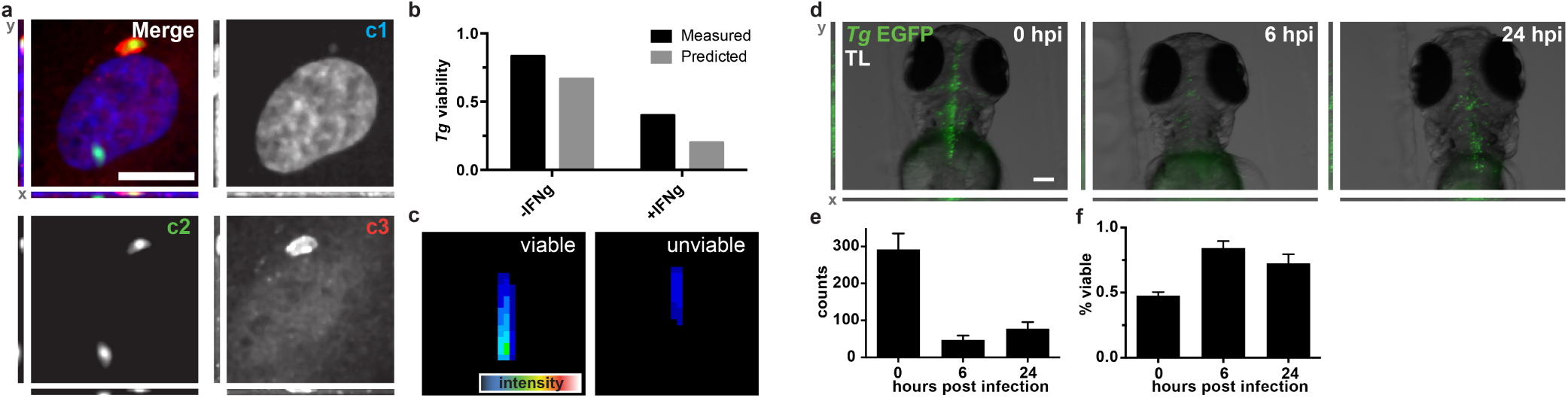
Mimicry embedding and weight transfer employed for *Toxoplasma gondii* (*Tg*) viability detection in cell culture and *in vivo*.. **a**, Merged three channel fluorescent image of a HUVEC cells infected *Tg* EGFP. Individual channels represent DNA stain (c1), *Tg* EGFP (c2), Ubiquitin (c3). Scale bar; 25 µm. **b**, Quantification of weak-labelled (measured) and CapsNet inferred (predicted) viable and unviable parasites. **c**, Representative reconstructions of the trained CapsNet network for viable and non-viable classes of *Tg* EGFP Z-profiles. **d**, Representative images (maximum intensity projections) of zebrafish (D. rerio) larvae infected with *Tg* EGFP at 0, 6 and 24 h pi. Scale bar; 100 µm. **e**, ZedMate detected *Tg* counts at 0,6 and 24h pi **f**, *In vivo* inference of *Tg* EGFP viability over time using DropConnect viability model trained on *in vitro Tg* data. N=10 images, error bars + SEM.

To assure the ANN could accurately distinguish between viable and unviable parasites we generated a data set of cells infected with *Tg*-EGFP using a specific viability label (c3) as ground truth (Fig.5a). To further assess viability, experiments were performed in the absence or presence of INFg, which drives parasite killing. Upon model training and validation, test inference on this dataset using c1-c2 resulted in a 84% and 80% match between measured (c3) and predicted (c1-c2) viability in the absence or presence of INFg, respectively (Fig.5b). CapsNet generator reconstructions showed that “viable” *Tg*-EGFP appear larger and brighter than “unviable” parasites in both c1 and c2 (Fig. 5c). This likely explains the ability of the model to accurately predict *Tg*-EGFP viability in the absence of specific c3 viability labelling.

In an attempt to train a general model for *in vivo* parasite viability assessment using our *in vitro* dataset, we performed mimicry embedding on *Tg*-EGFP (Fig. 5a; c2). This resulted in a >10% drop in prediction accuracy when training on CapsNet or using Autokeras (30)), a neural architecture search (data not shown). This suggested that single channel mimicry embedding does not provide enough context for training of complex algorithms. However, we reasoned as our mimicry embedding is based on MNIST, we could use any algorithm that performs well on this dataset. By switching to Drop-Connect (31) architecture, which performs among the best on MNIST, our classifier achieved 65% accuracy (precision 65.9%, recall 64.3%) in differentiating between viable and unviable parasites using a single channel (Fig. 1e-h for classifier training).

To test this classifier on an *in vivo* dataset we infected zebrafish (Danio rerio) larvae with *Tg*-EGFP and imaged them at 0, 6 and 24 h after infection by fluorescent 3D-stereomicroscopy (Fig. 5d). ZedMate was used to detect and quantify *Tg*-EGFP numbers over time (Fig. 5e). A dramatic drop-off in parasite count was seen between infection at 0 h and 6 h, followed by increased numbers of *Tg*-EGFP by 24h. Next the *Tg*-EGFP Z-profiles were mimicry embedded, normalized and their viability inferred using the *in vitro* infected cell model previously trained on DropConnect. At high pathogen load (0 h) 48% of *Tg*-EGFP were scored as viable (Fig. 5f). By 6 h this increased to 95% without any significant change within 24 h. These results are consistent with an initial clearing of unviable parasites, and replication of the remaining viable ones (32).

## Discussion

ANN analysis of biomedical datasets has trailed behind the unprecedented advancement of AI analysis seen in other fields. This is largely due to the lack of open source, verified biomedical datasets comparable to MNIST and ImageNet (19, 20). Here we present ZedMate and mimicry embedding as a strategy to harness the power of datasets like MNIST and transfer learning to train highly accurate models for analysis of 3-D biomedical data. ZedMate, an open source (ImageJ/Fiji) plugin designed for rapid detection and batch-quantification of 3D images at the single spot level made mimicry embedding possible.

When used together with CapsNet (15) mimicry embedding proved to be a promising method for detection of complex biomedical phenotypes *in vitro*. We show that transforming real-world images such that they resemble landmark datasets assures compatibility with, and seamless switching between, cutting-edge architectures. Embedding data in such a way allows one to maintain full compatibility with weights of the first layers thereby improving transfer. Using *in vivo* biomedical data, we further demonstrate that mimicry embedding can yield a model with higher accuracy than one obtained through cutting-edge neural architecture search. Thus, mimicry embedding can serve as a common denominator for assessing performance between architectures. Collectively, our results suggest that ZedMate and mimicry embedding, although employed here for the analysis of host-pathogen interaction, can be used for AI analysis of any biomedical 3-D dataset.

## Materials and Methods

### Cell culture, antibodies and reagents

HeLa cells (ATCC) were maintained in in Dulbecco’s modified Eagle’s medium (DMEM, Gibco, Life Technologies, Switzerland) with the addition of 10% fetal bovine serum (FBS, Sigma), and 1% penicillin-streptomycin (Pen-Strep, Sigma), 2 mM GlutaMAX (Life Technologies). Human Umbilical Vein Endothelial cells, HUVECs, (C12203, Promocell), were maintained in M199 medium (Gibco) supplemented with 30 mg/mL endothelial cell growth supplement (ECGS, 02–102, Upstate), 10 units/mL heparin (H-3149, Sigma) and 20% FBS (Sigma). Cells were cultivated on plates, pre-coated with 1% (w/v) porcine gelatin (G1890, Sigma). Both HUVECs and HeLa were grown as monolayers at 37.0°C and 5.0% CO2. HU-VEC were not used beyond passage 6.

Hoechst 33342 (Sigma) was used post fixation at 1:10,000 dilution throughout. Cell culture grade dimethyl sulfoxide (DMSO), used to dissolve control experimental compounds was obtained from Sigma.

### VACV and parapoxvirus strains and virus purification

Vaccinia virus strain Western Reserve expressing A5 mCherry protein (VACV WR) was used throughout (28, 33, 34). VACV mature virions (MVs) were purified from cytoplasmic lysates by being pelleted through a 36% sucrose cushion for 90 min at 18,000 × g. The virus pellet was resuspended in 10 mM Tris (pH 9.0) and subsequently banded on a 25 to 40% sucrose gradient at 14,000 × g for 45 min. Following centrifugation, the viral band was collected by aspiration and concentrated by pelleting at 14,000 × g for 45 min. MVs were resuspended in 1 mM Tris (pH 9.0), and the titter was determined for PFU per millilitre as previously described (35).

#### Early VACV infection and extracellular virions staining

HeLa cells were seeded onto CELLview Slide (Greiner Bio-One) at 10,000 cells per well 16h before the experiment. VACV A5-mCherry F13-EGFP was added at MOI 20, to increase the chances of synchronous infection. Cells were fixed with 4% EM- grade PFA 4 hours post infection (hpi) for 20 min followed by a PBS wash. Staining and labelling was preceded by blocking (without permeabilization) in blocking buffer (5% BSA, 1% FBS, in PBS) for 60 min at room temperature (RT). Next, L1 mouse (7D11) antibody (36) (1: 1000) in blocking buffer was added for 60 min at RT, followed by a PBS wash. Anti-mouse antibody (Alexa 647, Invitrogen. 1:1000), Phalloidin 594 (Sigma, 1:1000) and Hoechst in blocking buffer were added for 60 min at RT, followed by a PBS wash. 1,1’-Disulfanediyldinaphthalen-2-ol VACV entry inhibitor (IPA-3) was obtained and used as described (33). DMSO concentration was equal to or below 1%.

#### Late VACV infection and staining

HeLa cells we cultured on the coverslips and infected with VACV WR expressing A5 mCherry protein. At 8 hpi cells were fixed with 4% v/v FA. Next, VACV B5 protein antibody (mouse, 1:1000) in blocking buffer was added for 60 min at RT, followed by a PBS wash. Anti-mouse antibody (Alexa 647), Hoechst in blocking buffer were added for 60 min at RT, followed by a PBS wash.

#### *Toxoplasma gondii* (*Tg*) cultivation learning infection phenotypes

*Toxoplasma* (RH type I and Prugniaud type II strains) expressing GFP/luciferase were maintained *in vitro* by serial passage on human foreskin fibroblasts (HFFs) cultures (ATCC). Cultures were grown in DMEM high glucose (Life Technologies) supplemented with 10% FBS (Life Technologies) at 37°C in 5% CO_2_.

#### *Tg* cultured cells infection and staining

The day before the infection, type II parasites were passaged onto new HFFs to obtain parasites with a high viability. *Tg* were prepared from freshly 25G syringe-lysed HFF cultures. Parasites were subsequently 2 x 27G syringe lysed and excess HFF cell debris removed by centrifugation. Then, the parasites were added to the experimental cells at an MOI=2. The cell cultures with added *Tg* were then centrifuged at 500 x g for 5 min to synchronize the infection and the cultures incubated at 37°C in 5% CO_2_ for 3h. Samples treated with interferon gamma (IFN*γ*) were subjected to 100 IU/mL human IFN*γ* (285-IF, RD Systems) for 18h prior to infection. Upon fixation cells were stained with Hoechst 33342 and mouse mAb anti-ubiquitin FK2 (PW8810, Enzo Lifesciences; RRID: *AB*_1_0541840) and Alexa Fluor 568-conjugated secondary goat anti-mouse (A-11004, Invitrogen; RRID:*AB*_1_41371).

#### *Tg* infection *in vivo*

Tg EGFP parasites (type 1) were prepared from freshly 25G syringe-lysed HFF cultures in 10% FBS. Parasites were subsequently 27G syringe-lysed and excess HFF material removed by centrifugation. After washing with PBS, *Toxoplasma* tachyzoites were resuspended at 2.0×106 tachyzoites/µl in PBS.

Larvae were anesthetized with 20 µg/ml tricaine (Sigma-Aldrich) during the injection procedures and for all live *in vivo* imaging. All experiments were carried out on TraNac background larvae to minimize obstruction of fluorescence signal by pigmentation. 3dpf larvae were anesthetized and injected with 2.5 nl of parasite suspension into the hindbrain ventricle (HBV) as previously described (37). Infected larvae were transferred into individual wells containing 0.5x E2 media supplemented with methylene blue pre-warmed to 33°C.

#### Zebrafish husbandry and maintenance

Fish were maintained at 28.5°C on a 14hr light, 10hr dark cycle. Embryos obtained by natural spawning were maintained in 05x E2 media supplemented with 0.3 µg/ml methylene blue.

#### Ethics statement

Animal experiments were performed according to the Animals (Scientific Procedures) Act 1986 and approved by the Home Office (Project licenses: PPL P84A89400 and P4E664E3C). All experiments were conducted up to 4 days post fertilisation.

#### Super-resolution imaging of VACV intracellular virions

Supper-resolution microscopy was performed using a 100x oil immersion objective (NA 1.45) on a VT-iSIM microscope (Visitech; Nikon Eclipse TI), using 405 nm, 488 nm, 561 nm, 647 nm laser frequencies for excitation.

#### High-Content *Tg* EGFP imaging in cells

Black plastic flat-bottom 96-well plates (Falcon 353219) were imaged on an Opera Phenix High Content Imaging Platform using 63x magnification, 8 Z-slices (0.5 µm/slice) and multiple fields of view per well. Images were as single channel 16-bit tiff files and further processed for ZedMate analysis.

#### 3D *Tg* EGFP imaging *in vivo*

Progress of the *in vivo* infection was monitored by fluorescent stereomicroscopy (Leica M205FA, Leica Microsystems, Nussloch GmbH, Nussloch, Germany) at regular time points. All images were obtained with a 10x objective, at 13x magnification (0.79 µm/px) 20 z planes were captured covering a total distance of 171µm (8.55µm intervals).

#### Data processing and deep neural network training

Our training hardware was based on a single Nvidia 1080 Ti GPU set up in Intel Core i7 8700K system equipped with 32 Gb of RAM and an SSD. Installation consisted of Anaconda Python, Keras-gpu 2.2, Tensorflow-gpu 1.10 and KNIME 3.7.1. Some models were trained on 2019 MacBook Pro equipped with Intel Core i5 CPU using Keras 2.2 CPU. Source code is avalable under https://github.com/ayakimovich/ZedMate, example dataset under https://github.com/ayakimovich/virus-mnist. Further materials are available upon request.

## Appendix 1. Supplementary Information

This is a supplementary section to the preprint manuscript by Yakimovich et al. Source code is available under https://github.com/ayakimovich/ZedMate. All further information is available upon request.

**Figure S1.**
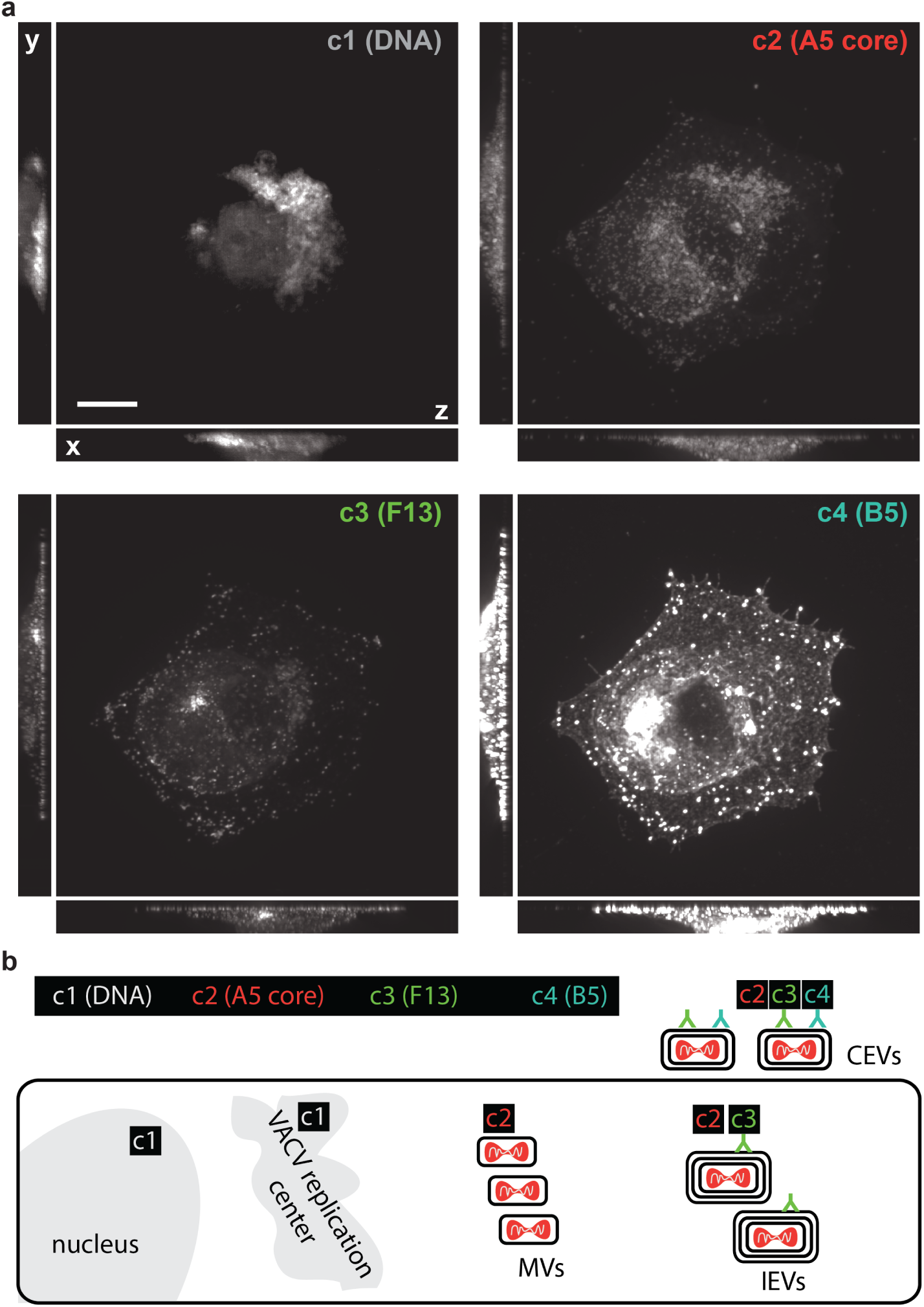
Individual channels used in late VACV infected cells and their biological relevance). **a**, Maximum intensity projections of individual channel images of HeLa cells infected with VACV at 8 hours post infection. Here, DNA stain (c1), VACV core A5-mCherry (c2), VACV outer envelope protein F13 (c3) and VACV outer envelope protein B5 (c4). Scale bar; 10 µm. **b**, Illustration of the position of markers in virions [MV (mature virions), IEV (intracellular enveloped virions), CEV (cell-associated extracellular virions)], and these virions - with the corresponding markers - in infected cells. Here c1 marks cellular DNA and cytoplasmic VACV replication sites, c2 marks all virions (MVs, IEVs and CEVs), c3 marks a subset of virions (IEVs and CEVs) and c4 marks only CEVs.

**Figure S2.**
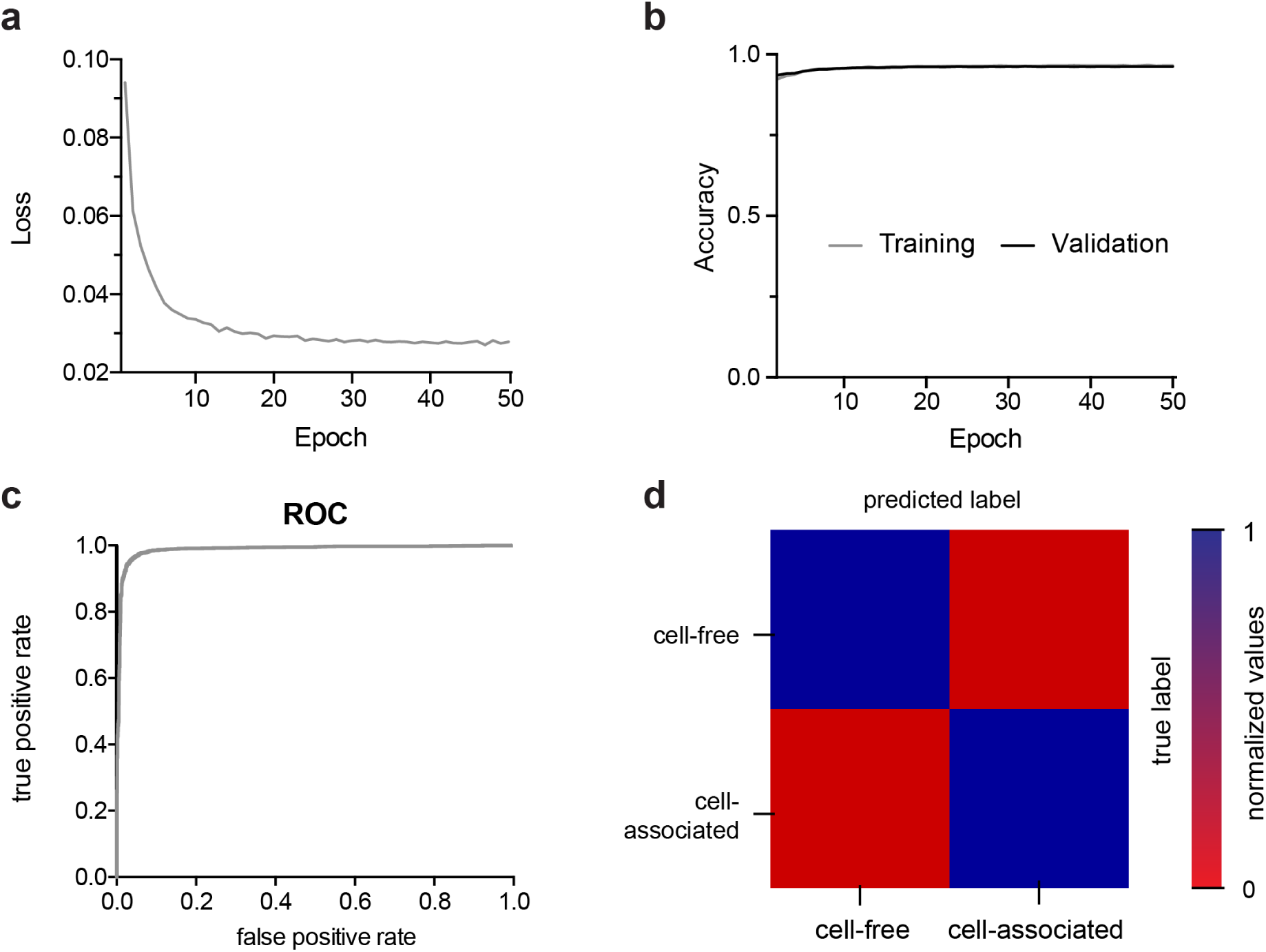
CapsNet training and validation of cell-free vs. cell-associated virus model. **a**, Late model loss function change upon training iterations (epochs). **b**, Late model training and validation (unseen data) accuracy change upon training iterations (epochs). **c**, Late model receiver operational characteristics (ROC) curve of the trained model obtained using unseen data (validation). Here area under the curve (AUC) was 0.989. **d**, Late model confusion matrix of the trained model obtained using unseen data (validation). Late model precision was 96.2%, recall was 96.2%, F1-score was 96.2%.

**Figure S3.**
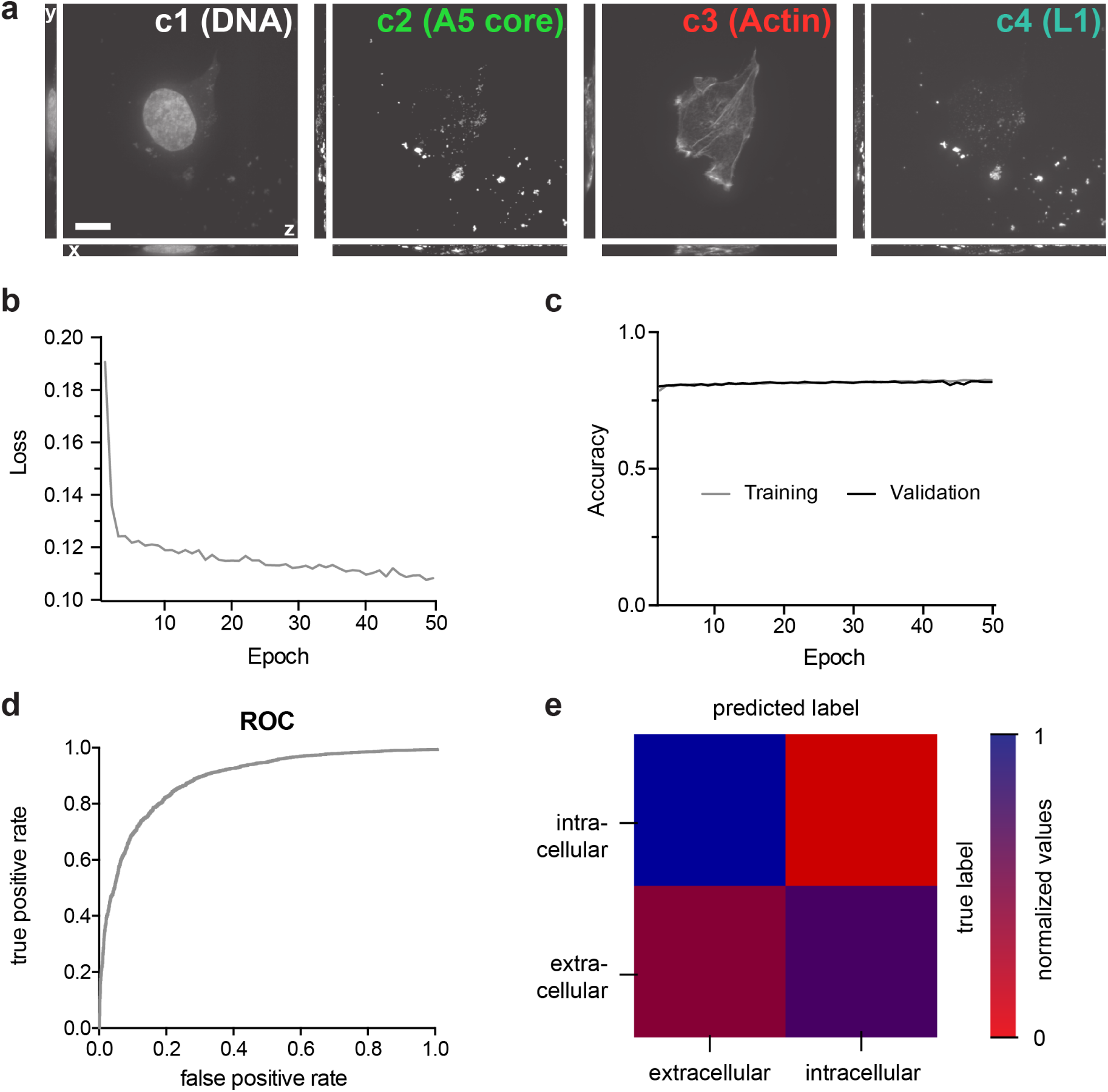
CapsNet training and validation of extracellular vs. intracellular virus model. **a**, Maximum intensity projections of individual channels from HeLa cells infected with VACV. Here, DNA stain (c1), VACV core A5-EGFP (c2), actin stained with phalloidin (c3) and VACV membrane protein L1 as an extracellular virion label (c4) **b**, Model loss function change upon training iterations (epochs). **c**, Model training and validation (unseen data) accuracy change upon training iterations (epochs). **d**, Model receiver operational characteristics (ROC) curve of the trained model obtained using unseen data (validation). Here area under the curve (AUC) was 0.896. **e**, Model confusion matrix of the trained model obtained using unseen data (validation). Model precision was 81.3%, recall was 81.4%, F1-score was 81.8%.

**Figure S4.**
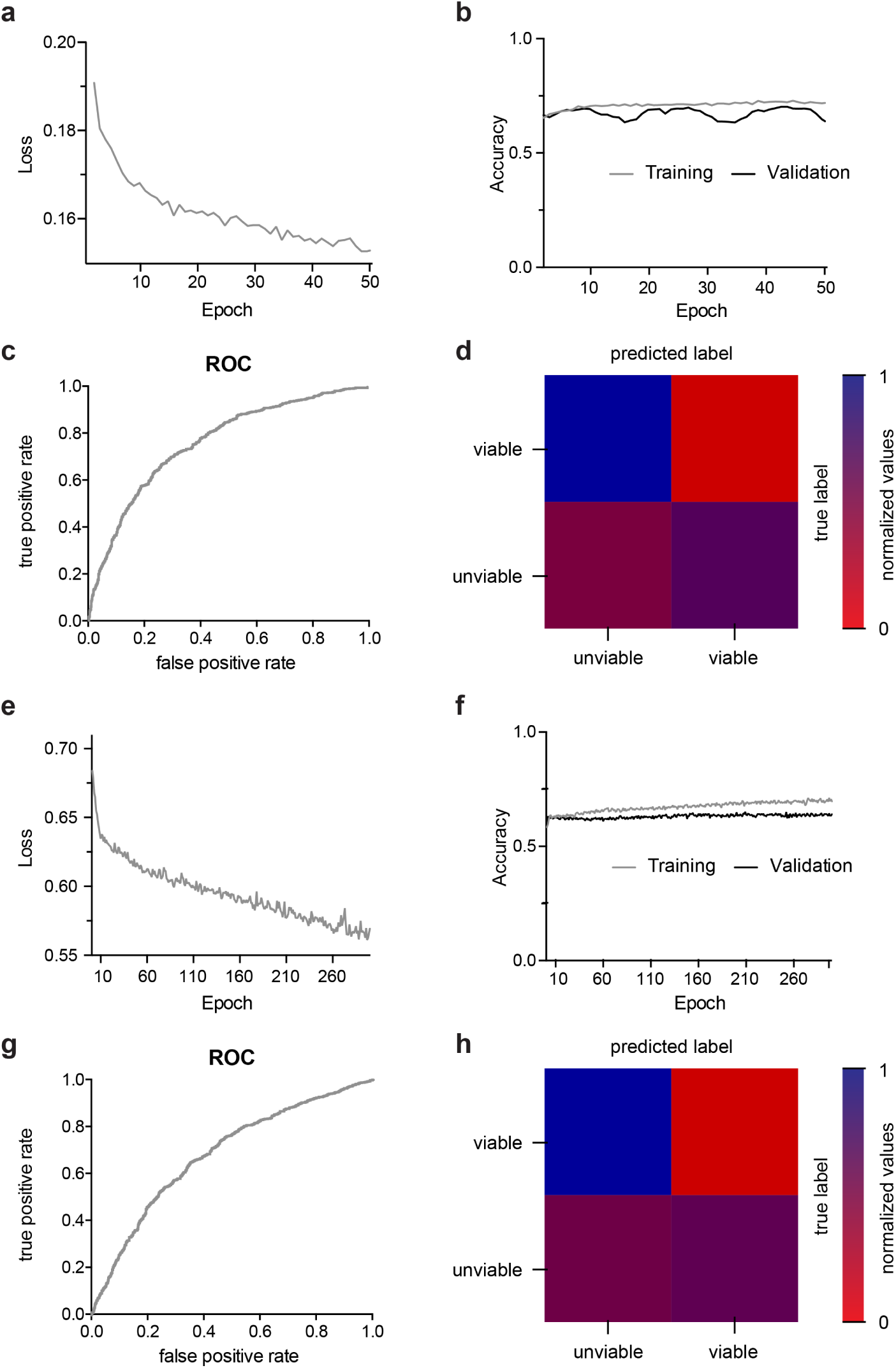
CapsNet and DropConnect training and validation of *in vitro* and *in vivo Tg* EGFP viability model. **a**, 2-channel CapsNet model loss function change upon training iterations (epochs). **b**, 2-channel CapsNet model training and validation (unseen data) accuracy change upon training iterations (epochs). **c**, 2-channel CapsNet model receiver operational characteristics (ROC) curve of the trained model obtained using unseen data (validation). Here area under the curve (AUC) was 0.764. **d**, 2-channel CapsNet model confusion matrix of the trained model obtained using unseen data (validation). **e**, 1-channel DropConnect model loss function change upon training iterations (epochs). **f**, 1-channel DropConnect model training and validation (unseen data) accuracy change upon training iterations (epochs). **g**, 1-channel DropConnect model receiver operational characteristics (ROC) curve of the trained model obtained using unseen data (validation). Here area under the curve (AUC) was 0.685. **g**, 1-channel DropConnect model confusion matrix of the trained model obtained using unseen data (validation). The 2-channel CapsNet model precision was 72.5%, recall was 70.7%, F1-score was 70.1%. The 1-channel DropConnect model precision was 65.9%, recall was 64.3%, F1-score was 64.9%.

